# Mitochondrial DNA Copy Number Variation Across Human Cancers

**DOI:** 10.1101/021535

**Authors:** Ed Reznik, Martin L. Miller, Yasin Senbabaoglu, Nadeem Riaz, William Lee, Chris Sander

## Abstract

In cancer, mitochondrial dysfunction, through mutations, deletions, and changes in copy number of mitochondrial DNA (mtDNA), contributes to the malignant transformation and progress of tumors. Here, we report the first large-scale survey of mtDNA copy number variation across 21 distinct solid tumor types, examining over 13,000 tissue samples profiled with next-generation sequencing methods. We find a tendency for cancers, especially of the bladder and kidney, to be significantly depleted of mtDNA, relative to matched normal tissue. We show that mtDNA copy number is correlated to the expression of mitochondrially-localized metabolic pathways, suggesting that mtDNA copy number variation reflect gross changes in mitochondrial metabolic activity. Finally, we identify a subset of tumor-type-specific somatic alterations, including IDH1 and NF1 mutations in gliomas, whose incidence is strongly correlated to mtDNA copy number. Our findings suggest that modulation of mtDNA copy number may play a role in the pathology of cancer.

## 1 Introduction

Human cells contain many copies of the 16 kilobase mitochondrial genome, which encodes thirteen essential components of the mitochondrial electron transport chain. Alterations of mitochondrial DNA (mtDNA), via inactivating genetic mutations or depletion of the number of copies of mtDNA in a cell, impair mitochondrial respiration and contribute to pathologies as diverse as encephelopathies and neuropathies [9], and the process of aging [1,11]. In cancer, a number of studies have examined the role of mtDNA mutations in carcinogenesis [13, 16, 19, 41]. However, the contribution of changes in the gross number of mtDNA molecules in a tumor (*i.e*. the “mtDNA copy number”) to tumor development and progression has not been adequately investigated.

In contrast to the fixed (diploid) copy number of the nuclear genome, many copies of mtDNA exist within each cell, and these levels can fluctuate based on energetic demand. Because mitochondria undergo a constant process of fusion and fission, it is difficult to meaningfully determine the number of mtDNA molecules per mitochondrion. Instead, studies have focused on measuring mtDNA copy number per cell, with estimates that vary between a few hundred and over one hundred thousand copies, depending on the tissue under examination [40]. Furthermore, because mtDNA serves as a template for the transcription of essential electron transport chain complexes, the quantity of mtDNA in a cell is a surrogate marker for the cell’s capacity to conduct oxidative phosphorylation. For instance, a recent study estimated that energy-intensive tissues such as cardiac and skeletal muscle contained between 4000 and 6000 thousand copies of mtDNA per cell, while liver, kidney, and lung tissues averaged between 500 and 2000 copies [8].

Mitochondrial dysfunction plays several distinct roles in cancer [19, 34, 41]. First, the normal functions of mitochondria (*e.g.* respiration) may be subverted to support the growth of the tumor. A canonical example of this is the observation that many tumors suppress mitochondrial respiration in favor of increased uptake of glucose and secretion of lactate (“the Warburg effect”), a phenomenon which has found clinical utility for imaging of tumors using FDG-PET [39]. Second, mitochondria are susceptible to mutations in nuclear-and mitochondrially-encoded genes, and a subset of tumors are known to be caused by mutations of the mitochondrial enzymes FH,SDH, and IDH [18]. Furthermore, mtDNA dysfunction affecting the electron transport chain can lead to generation of excess reactive oxygen species (ROS), contributing to tumor cell metastasis [15].

To date, no comprehensive analysis of mtDNA copy number changes in tumors has been completed, despite a large literature of isolated reports [46]. Large-scale studies of mtDNA in cancer have instead focused on the analysis of mutations and heteroplasmy, largely ignoring the contribution of mtDNA copy number variation to the development and progression of tumors. Here, we use whole-genome and whole-exome sequencing data to examine changes in mtDNA copy number across a large panel of cancer types profiled by The Cancer Genome Atlas (TCGA) consortium. Using the resulting mtDNA copy number estimates, we ask fundamental questions about mtDNA and cancer. We investigate whether evidence of the Warburg effect can be found in patterns of mtDNA accumulation or depletion, which reflect a tumor’s capacity to respire. We examine the connection between gene expression levels and mtDNA copy number, and identify a subset of mitochondrially-localized metabolic pathways exhibiting a high degree of co-variation with mtDNA levels. Finally, we ask whether gross variations of mtDNA copy number are linked to the incidence of somatic alterations (including mutations and copy number alterations) across cancer types. Altogether, our results shed light on the contribution of aberrant mitochondrial function, through changes in mtDNA content, to cancer.

**Figure. 1.**
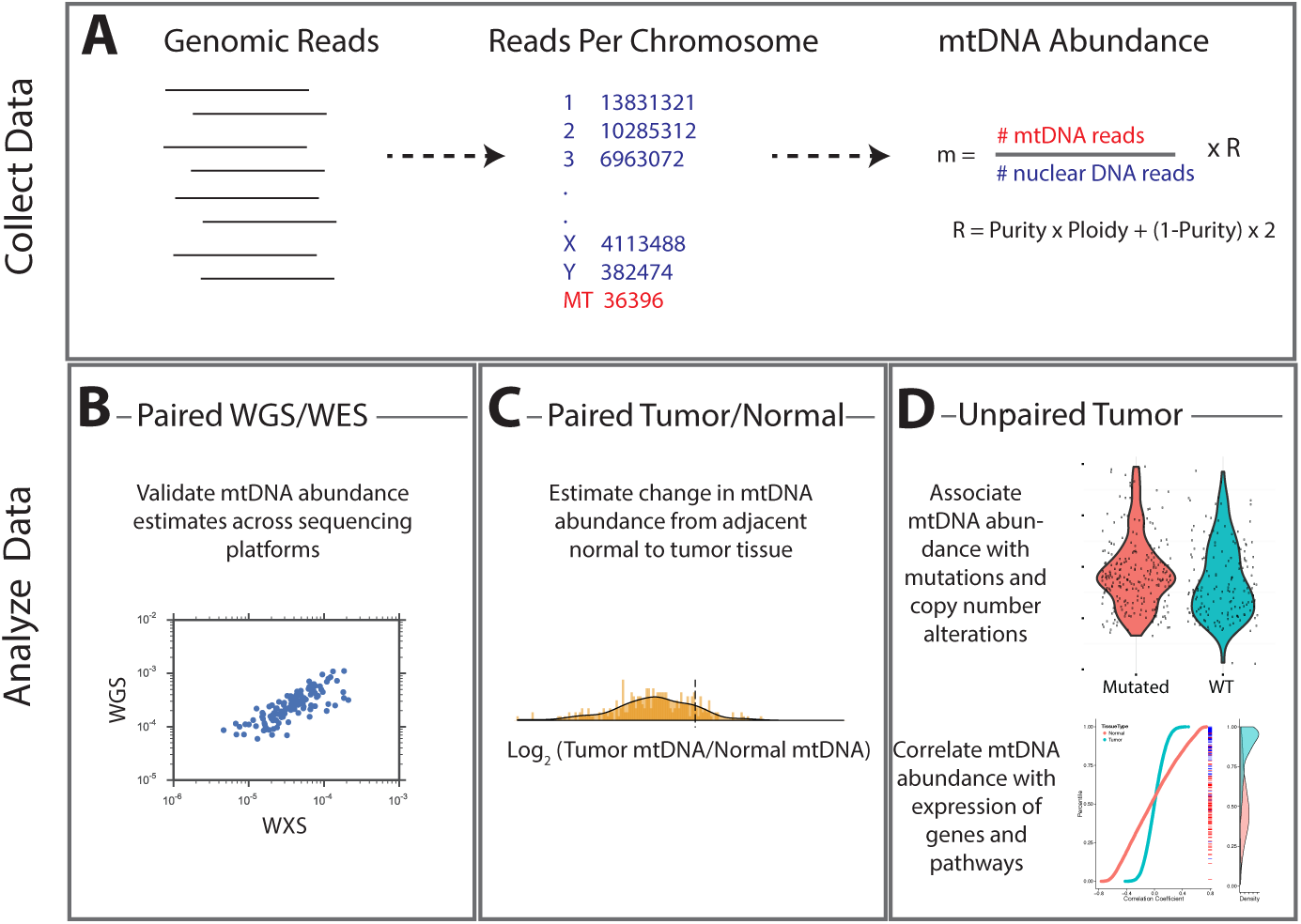
(A) Aligned reads are analyzed to determine the number of primary, properly paired reads aligning to each chromosome. Relative abundance of mitochondrial DNA is calculated as the ratio of mtDNA reads to nuclear DNA reads, and corrected for tumor purity and ploidy. The results of these calculations were employed in three different types of analysis. (B) Comparisons across samples profiled by both whole exome and whole genome sequencing provided validation of mtDNA copy number estimates. (C) Pairs of matched tumor/adjacent-normal samples were compared to uncover patterns of mtDNA accumulation and depletion. (D) Using all data available (including tumor samples lacking matched normal samples), statistical associations between mtDNA copy number and (1) mutation/copy number alterations, and (2) gene expression, were calculated.

## 2 Calculation of mtDNA Abundance

To estimate the copy number of mtDNA in a tumor sample, we implemented a computationally efficient and fast approach based on comparing the number of sequencing reads aligning to (1) the mitochondrial (MT) genome and (2) the nuclear genome. Comparable approaches have been used to estimate somatic copy number alterations within the nuclear genome in cancer (for a review, see [47]). The approach assumes that regions of the genome of equal ploidy should be sequenced to comparable depth. In a normal human cell, the autosomal nuclear genome is at a fixed (diploid) copy number. Thus, by calculating the ratio of reads aligning to the mitochondrial and nuclear genomes, respectively, it is possible to estimate mtDNA ploidy relative to a diploid standard. This approach to assaying mtDNA copy number has been proposed and implemented by others in prior work [8, 12, 32].

To estimate mtDNA copy number, we calculated the ratio of (1) the number of sequencing reads mapping to the MT genome (*r_m_*) to (2) the number of reads mapping to the nuclear genome (*r_n_*) (Equation 1). Because tumor cells can exhibit large-scale amplifications and deletions, and be infilitrated by stromal and immune cells, we applied a ploidy/purity correction (“*R*”), described in detail in the Methods. This calculation yields the relative mtDNA copy number *m*. Assuming two samples have been processed in identical manners, then the sample with a higher value of *m* contains more copies of mtDNA [8, 12]. Because the approach relies simply on counting the number of reads aligning to each chromosome and does not calculate statistics at the individual-exon level, the running time was fast (on the order of a few minutes per sequencing sample) and the calculations easily parallelizable across the thousands of samples interrogated.

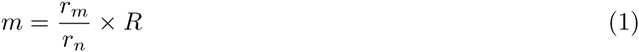

We applied this method to whole exome sequencing (WXS) and whole genome sequencing (WGS) data from 21 distinct TCGA studies (Figure 2, see Methods for further details on data collection). To validate estimates of mtDNA copy number, we compared estimates from samples submitted to both WXS and WGS. Although mitochondrial reads are abundant in both WGS and WXS, the two sequencing methods capture mtDNA at different efficiencies: exome sequencing involves the targeted enrichment of exonic regions prior to sequencing and does not target mtDNA [31], while WGS sequences cellular DNA in an unbiased manner. If our approach to estimating mtDNA copy number is accurate, then we expect that the two sequencing platforms should offer comparable estimates of mtDNA copy number across a panel of samples,*i.e*. samples with high mtDNA copy number in WGS should have similarly high mtDNA copy number in WXS. We compared mtDNA copy number estimates in 1174 samples across 9 tumor types profiled by both WXS and WGS, controlling for sequencing center due to previously reported biases in exome enrichment across different centers [16]. We confirmed that across all combinations of cancer types and sequencing centers, WXS and WGS offer significantly correlated estimates of mtDNA copy number (Table 1 and in the SI Figure S1).

**Figure. 2.**
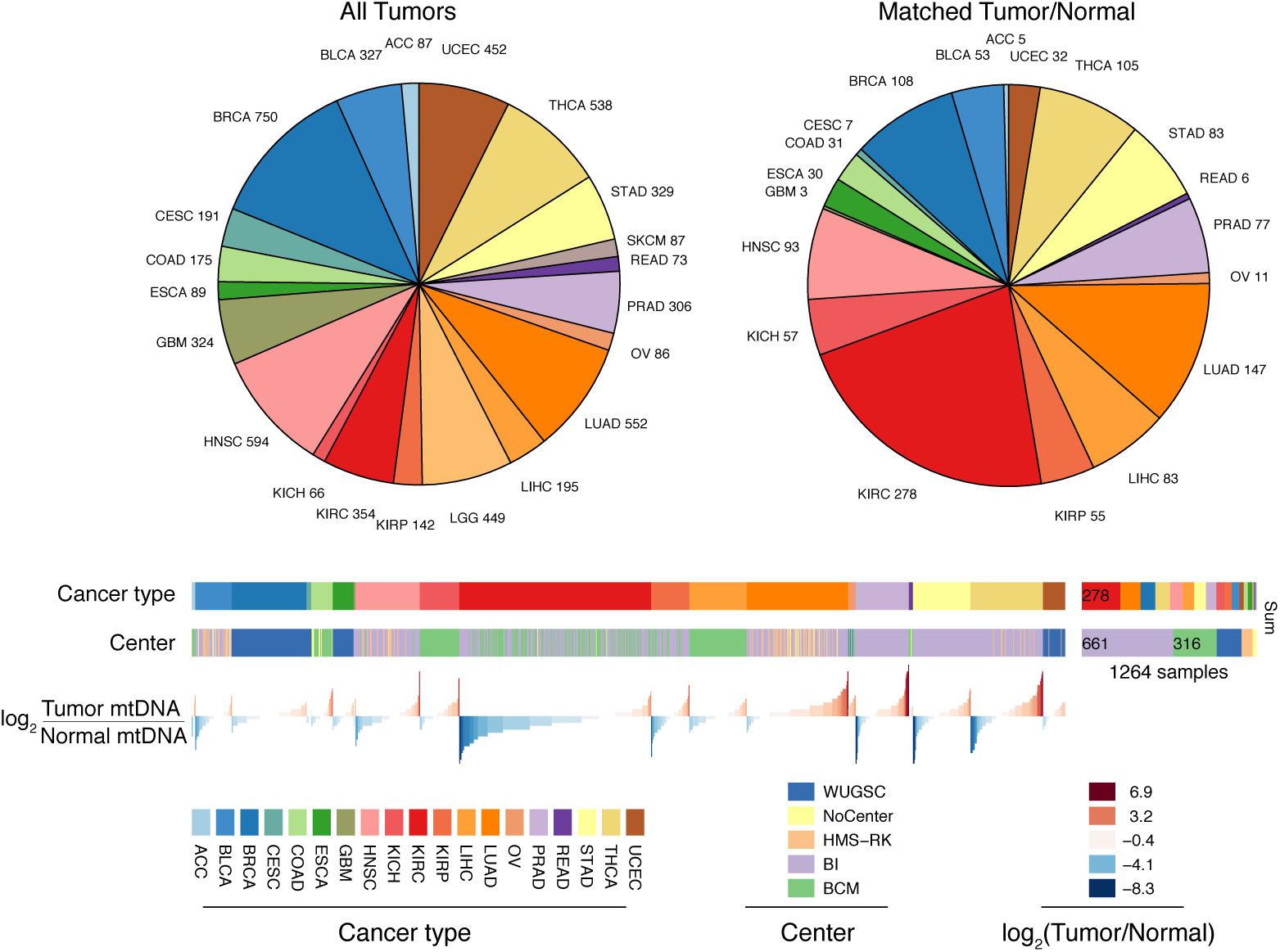
Summary of Data. Whole-exome and whole-genome sequencing data was obtained from 21 TCGA studies. The data was processed at 5 different sequencing centers, each of which was analyzed separately. Over 1400 of our samples are paired instances of tumor/adjacent-normal tissue from the same patient, which are used to quantify changes in mtDNA content across tumors.

## 3 Gross Changes in mtDNA Content are Evident in Many Cancers

Do tumors harbor different numbers of copies of mtDNA than normal tissue? This question is particularly intriguing because many cancers divert metabolic flux away from mitochondrial respiration and towards glycolysis in a phenomenon termed the Warburg effect. We investigated if tumor samples showed a significant depletion in mtDNA content, relative to matched normal tissues. To do so, for each pair of tumor/adjacent-normal samples collected from the same patient (1264 patients in total), we calculated the ratio *r* between the mtDNA copy number *m* in each of the samples:

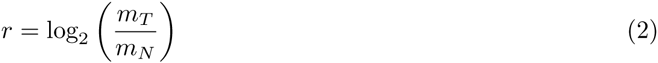

where *m_T_* and *m_N_* are the mtDNA abundances in tumor and normal tissues, respectively. We then used non-parametric Wilcoxon signed rank tests to assess whether each cancer type was signficantly enriched for tumor samples with higher or lower mtDNA than matched normal tissue. The analysis was restricted to 14 cancer types for which we had at least 10 matched tumor/normal pairs. To ensure a meaningful comparison, we only used adjacent-normal tissue (and not blood) for the analysis. We elected to focus on analyzing whole-exome sequencing data, for which we had the largest number of samples. A complete list of all results is available in Supplementary Table 2.

Strikingly, eight of the fourteen tumor types analyzed showed a statistically significant (Bonferonni-corrected Mann-Whitney p-value < 0.05) decrease in mtDNA abundance in tumor samples (Figure 3A). These “mtDNA-depleted” tumor types included bladder, breast, esophogeal, head and neck squamous, kidney (both clear-cell and papillary subtypes), liver, and stomach cancers. Despite a tendency towards mtDNA depletion, all tumor types contained at least one sample with higher mtDNA content than adjacent normal tissue. Nevertheless, the depletion effect was exceptionally strong in several tumor types: over 90% of bladder tumor samples and 88% of clear-cell kidney tumor samples contained less mtDNA than their normal tissue counterparts. In contrast, a single tumor type, lung adenocarcinoma, showed statistically significant mtDNA accumulation. In cases where sufficient numbers of both WGS and WXS data were available (bladder, breast, head and neck, clear-cell kidney, and lung adenocarcinomas), we observed a consistent effect across samples processed by both platforms (Figure S2).

**Figure. 3.**
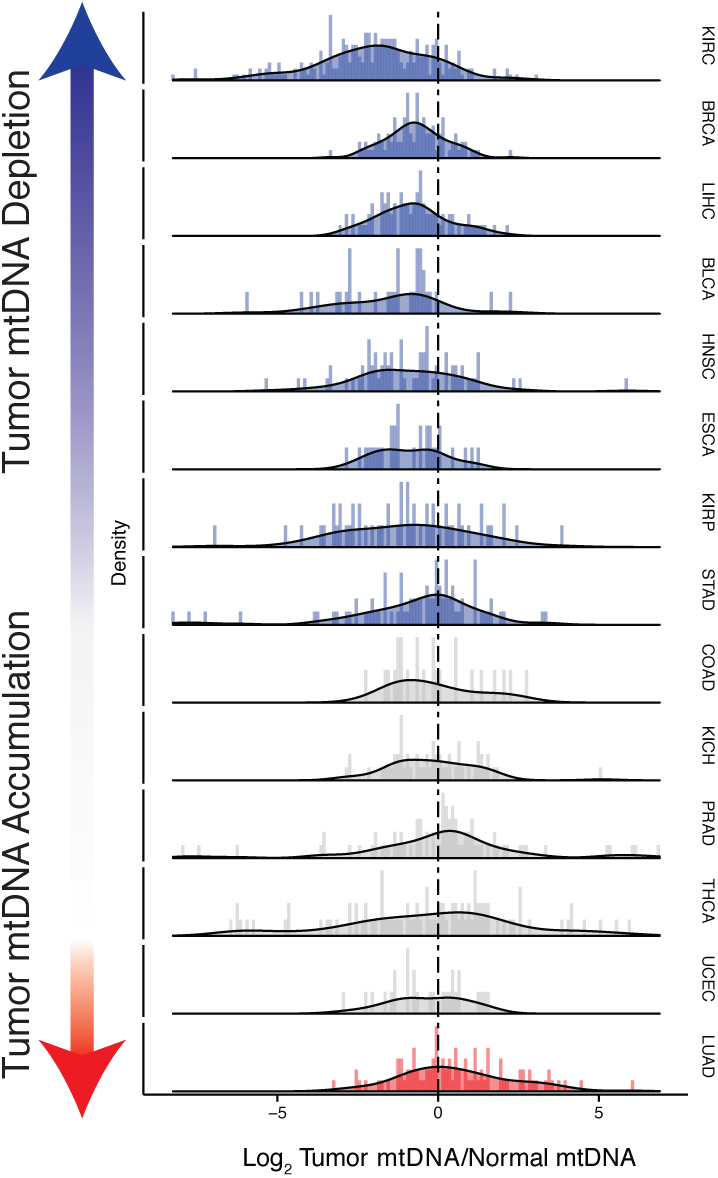
Many tumor types show depletion of mtDNA in tumor samples, relative to adjacent normal tissue. Normalized histograms and density plots illustrate log_2_ ratio of mtDNA content in tumor tissue, to mtDNA content in normal tissue. Tumor types with significant changes in mtDNA copy number changes are indicated in the sidebar with colored circles. Statistical significance of trends is assessed using Wilcoxon rank sign, with -logio p-values reported in the sidebar by the radius of the circle.

To further relate changes in mtDNA content to clinical progression of disease, we used Cox regression to determine if tumor mtDNA copy number was a significant predictor of patient survival (Figure 4). In total, 5 cancer types showed statistically significant association between patient survival and mtDNA content. In three cancer types (ACC, p-value 0.017;KICH, p-value 0.058; and LGG, p-value 0.007), high tumor mtDNA content was associated with better survival. The opposite trend, of poor survival in patients with high tumor mtDNA, was observed in SKCM (p-value 0.038), and UCEC (p-value 0.004). The finding regarding KICH is particularly intriguing given the central role mitochondrial dysfunction has been proposed to play in the disease [7]. That mtDNA copy number correlates with better or worse survival, depending on cancer type, suggests that other confounding factors strongly tied to survival, such as the presence of somatic mutations, may influence mtDNA levels. In a later section, we will investigate this hypothesis.

**Figure. 4.**
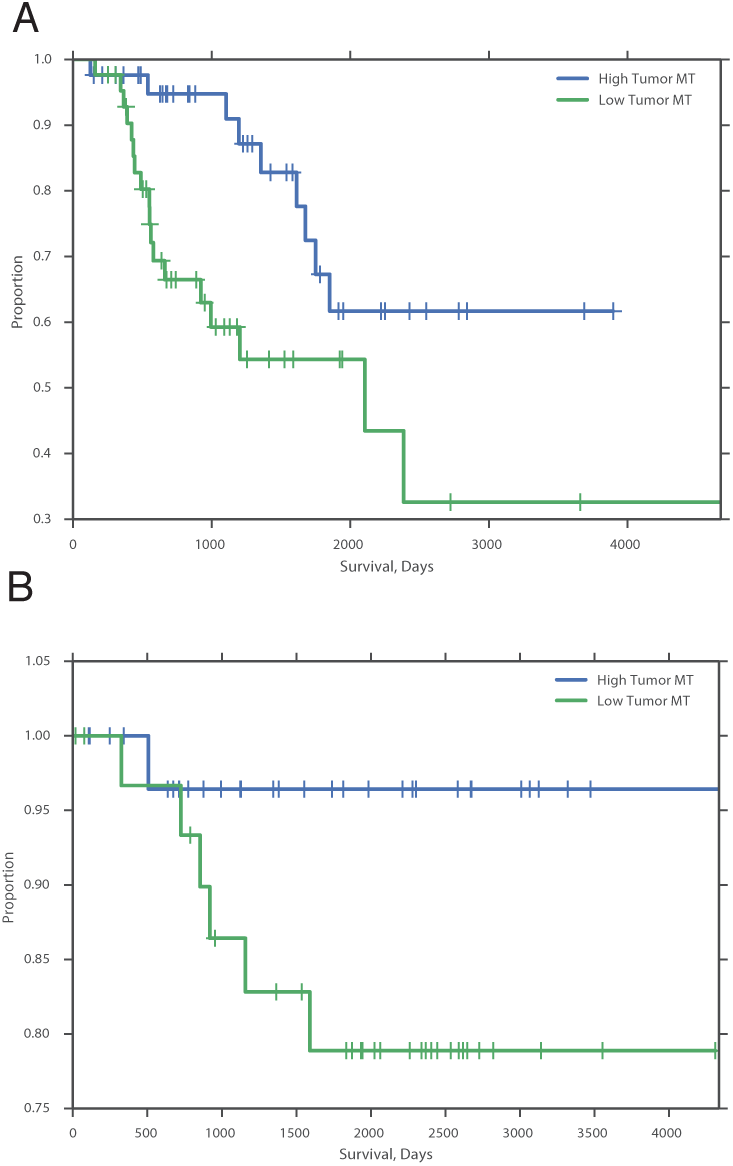
mtDNA content is significantly associated with patient survival in (A) adrenocortical (ACC) and (B) kidney chromophobe carcinoma (KICH). For visualization purposes, patients are partitioned into two groups, based on tumor mtDNA copy number relative to the median mtDNA copy number across all tumor samples in the cancer type. Cox regression identified a significant association between high tumor mtDNA and better survival (ACC, p-value 0.017; KICH, p-value 0.058).

## 4 mtDNA is Correlated to the Expression of Mitochondrial Metabolic Genes

Proteins encoded in mtDNA localize exclusively to the mitochondrial electron transport chain, and fluctuations in mtDNA copy number are well-known to influence the level of transcription of these genes. It has also been observed that complete depletion of mtDNA in cell lines by exposure to ethidium bromide affects a number of additional signaling pathways [2]. Thus, we were compelled to ask if changes in mtDNA content narrowly influenced changes in the expression of oxidative phosphorylation genes, or if they were more broadly connected to the other functions of mitochondria.

Our approach to this question was to search for gene sets whose transcriptional signatures were highly correlated to mtDNA copy number. To do so, we calculated the non-parametric Spearman correlation between each gene and mtDNA copy number, and then used the mean-rank gene set test implemented in limma [20] to identify gene sets which were significantly enriched for highly correlated genes. The approach was applied in an unbiased manner to all 674 Reactome gene sets in the Canonical Pathways group from the MSigDB database [24].

In general, each tissue exhibited specific gene sets which were strongly correlated to mtDNA copy number levels. However, when aggregating across all cancer types, mitochondrially-localized metabolic pathways showed the highest correlation with mtDNA abundance (Figure 5). The recurrent positive correlation between expression of mitochondrial genes and mtDNA copy number across many tumor types served as a second, independent validation that estimates of mtDNA copy number reflected *in vivo* mtDNA ploidy.

In line with expectation, we found that the “Respiratory Electron Transport” gene set was the most correlated to mtDNA copy number (1st out of 674 gene sets, Supplementary Table 3, Worksheet Fig5Data). Among the remaining top positively correlated gene sets, all were metabolism-related, and included the TCA cycle, mitochondrial beta oxidation of fatty acids, and branched chain amino acid (BCAA) catabolism. BCAAs (valine, leucine, isoleucine) are essential amino acids whose catabolism depends on the activity of an enzyme complex, branched chain *alpha-keto* acid dehydrogenase, located in the inner mitochondrial membrane [14], and prior work has demonstrated that dietary supplementation of BCAA’s promotes mitochondrial biogenesis [6, 37]. Furthermore, a recent study has shown that elevated plasma levels of BCAAs are found 2 to 5 years before a cohort of patients developed pancreatic ductal adenocarcinoma [26].

A number of gene sets also showed recurrent negative correlation to mtDNA copy number, including those related to mRNA splicing, and S-phase (2nd and 5th most anticorrelated out of 674 gene sets, Supplementary Table 3). Several of these gene sets are associated with known non-metabolic functions of mitochondria in the cell. In particular, the replication of mitochondria and mtDNA is intimately linked to the cell cycle [3], and the nucleotide precursors to mtDNA are in part produced *de novo,* via a pathway that is only active during the S phase of the cell cycle [35]. Interestingly, we did not observe a strong correlation or anti-correlation between mtDNA copy number and glycolysis, suggesting that the transcriptional regulation of respiration and glycolysis may be independently coordinated.

A subset of tumor types did not show strong positive correlation between mtDNA copy number and expression of mitochondrial metabolic genes. In some cases, this was the result of an apparently dominant correlation with another pathway. For example, in UCEC, there is an exceptionally strong correlation between expression of cell cycle genes and mtDNA copy number. Perhaps most interestingly, in both prostate tumors and adjacent normal tissue, the expression of mitochondrial respiratory genes was the most strongly anti-correlated gene set (see SI Table 3). We speculate that this effect may be associated with the unique mitochondrial metabolism of prostate epithelia, which secrete large amounts of citrate generated in the mitochondria, rather than oxidizing it further and using the resulting NADH in the respiratory electron transport chain [4, 5].

**Figure. 5.**
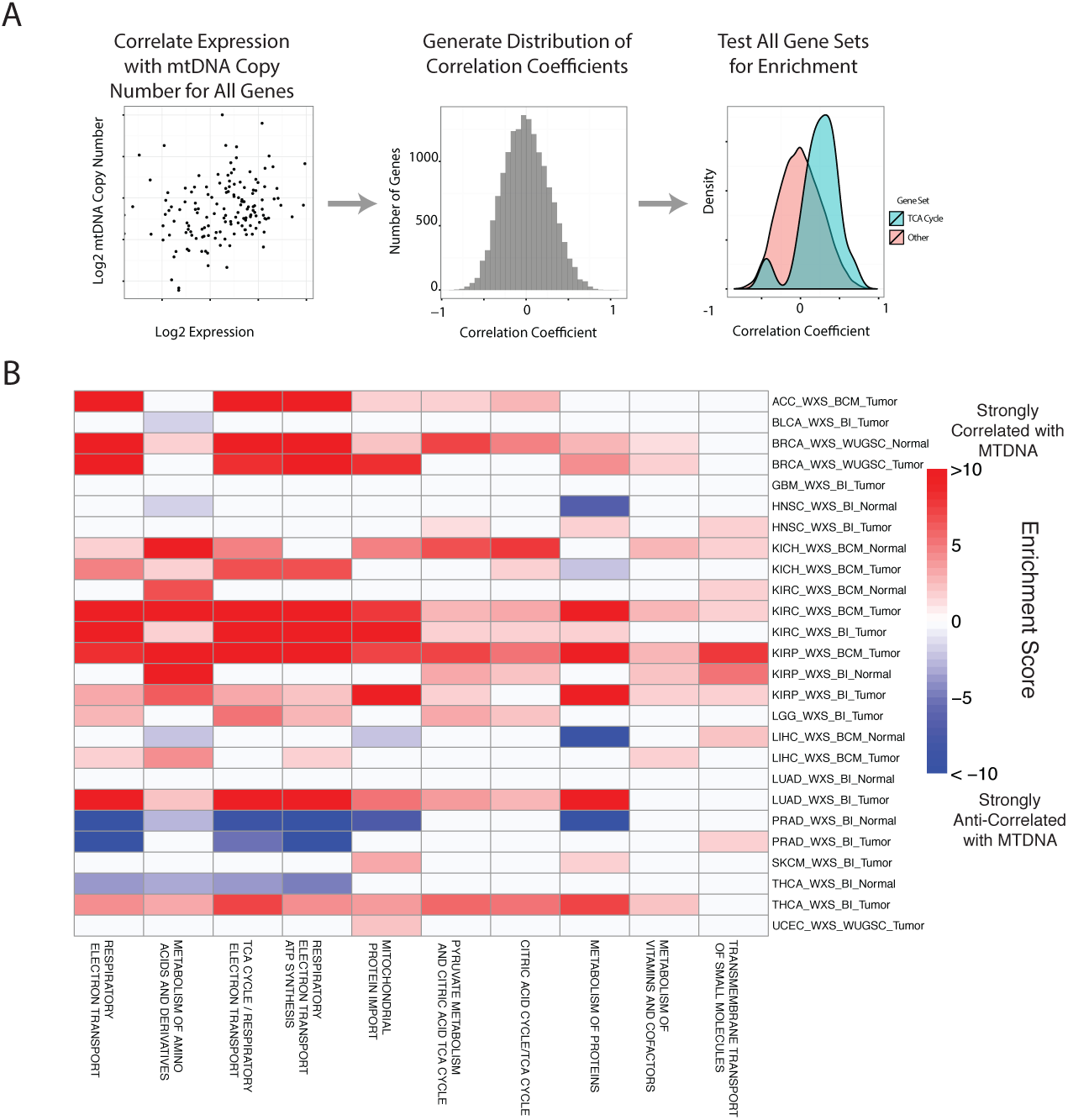
Gene set analysis identifies pathways correlated to mtDNA content. (A) Correlations between each gene and mtDNA content is calculated. Then, gene sets enriched for high/low correlation coefficients are identified. (B) mtDNA copy number is most strongly correlated to metabolic pathways including respiratory electron transport and the TCA cycle, which are localized to the mitochondria.

## 5 Association with Mutations and Copy Number Alterations

The landscape of genetic events driving tumors is diverse, and the presence and activity of these genetic lesions is now being used as the basis for the staging of tumors, design of clinical trials, and development of new treatments [30]. We sought to understand whether mtDNA abundance was associated with the incidence of particular mutations/copy number alterations (CNAs) in patient samples. To do so, we evaluated whether patients harboring a particular genetic lesion showed statistically significant increases or decreases in tumor mtDNA abundance. We restricted analysis to whole-exome sequencing data, using tumor types which were sequenced at a single sequencing center, and which were not under embargo by the TCGA as of March 2015. All results for the analysis are reported in Figure 6 and Supplementary Tables 4 and 5.

The most apparent result of our analysis was the association of a large number of CNAs in endometrial carcinomas (UCEC) with increased mtDNA abundance. Recent work by the TCGA proposed a subtype stratification of endometrial carcinomas based on mutation and CNA frequency [17]. Among these subtypes is a serous-like “copy-number-high” subtype with large numbers of somatic CNAs and extremely poor survival. We obtained the UCEC subtype classifications and confirmed that serous-like endometrial carcinomas exhibited substantially higher mtDNA copy number than all other subtypes (Mann-Whitney p-value 3 × 10^−10^, Figure 6), explaining the large number of associations we observed. A number of genes are also known to be enriched or depleted for mutations in the serous-like subtype, and these mutations also showed statistically significant association with mtDNA abundance (enriched: *TP53,* BH-corrected p-value 4 × 10^−5^, depleted: *PTEN,* BH-corrected p-value 4 × 10^−5^). To our knowledge, this is the first report of an association between the serous-like subtype of endometrial carcinoma and mtDNA copy number.

**Figure. 6.**
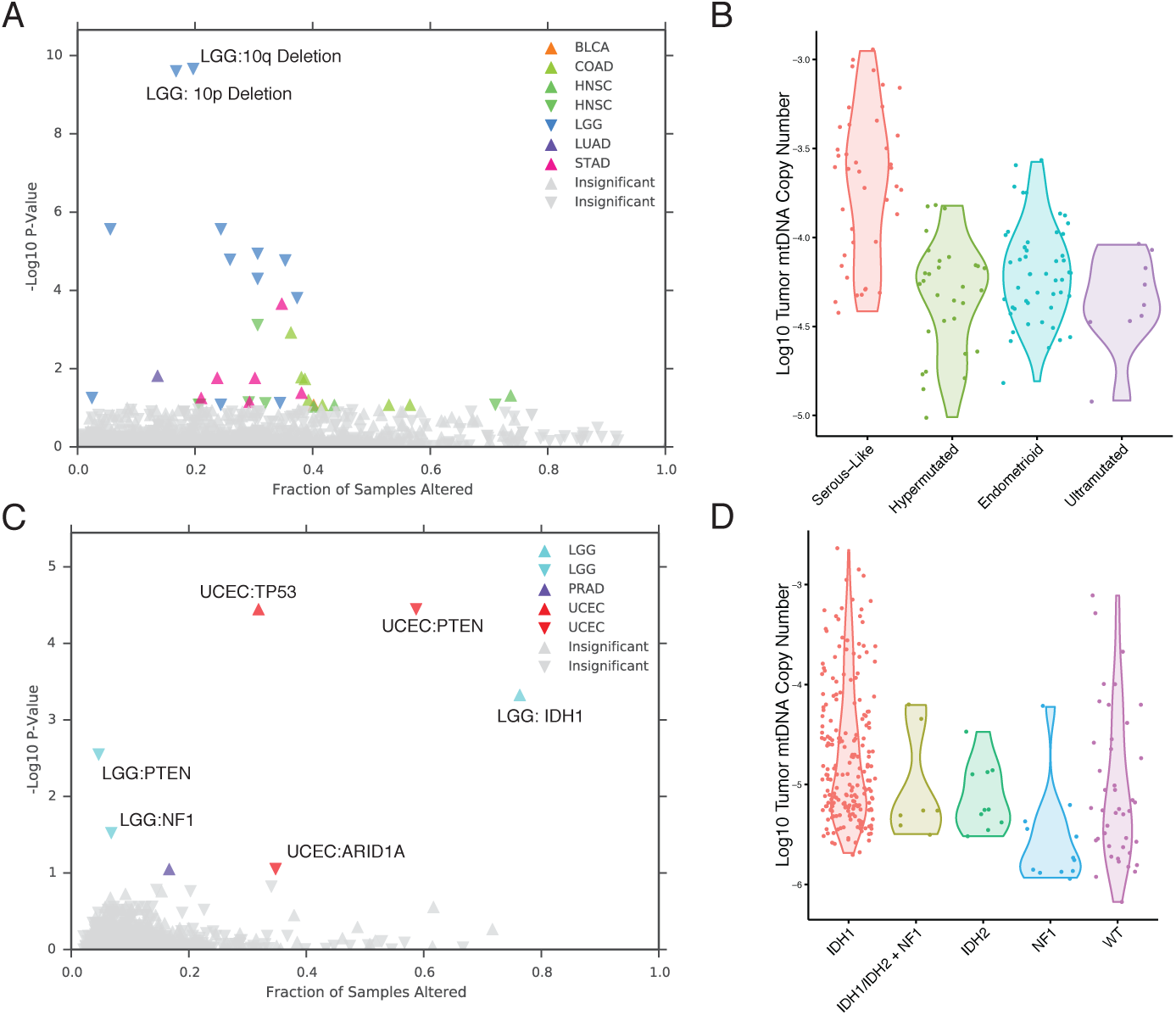
mtDNA content is correlated to the incidence of certain mutations and copy number alterations. Each point corresponds to a single alteration (e.g *P53* mutation). Direction of arrow indicates whether alteration increases or decreases mtDNA content. X-axis in (A) and (B) indicates the fraction of samples in a cancer-type that contained the alteration (*i.e*. ≈ 25% of LGG samples were 10q deleted). (A) 31 out 1434 CNAs tested were found to be significantly associated with mtDNA content. CNAs from the UCEC cancer type were filtered out because of strong association between mtDNA copy number and the “copy-number high” serous-like subtype in UCEC, shown in (B). (B) The UCEC serous-like subtype displays a marked increase in tumor mtDNA copy number, relative to tumors of other subtypes. (C) Relatively few mutations (7 out of 1766 tested) associate significantly with tumor mtDNA content. The UCEC associations are likely the result of the correlation to the serous-like subtype. (D) *IDH1* and *NF1* mutation status is strongly linked to increased tumor mtDNA copy number in LGG.

After filtering out associations in UCEC, we were left with a small number of statistically significant mutations and CNAs associated with mtDNA abundance. Among these, the strongest signal arose from increased tumor mtDNA content in *IDH1*-mutant low grade gliomas (Figure 6, BH-corrected p-value 5 × 10^−4^). Both *IDH1* and *IDH2* activating mutations induce production of the so-called “onco-metabolite” 2-hydroxyglutarate, which competitively inhibits α-ketoglutarate-dependent histone demethylases and 5-methylcytosine hydroxylases, inducing a hypermethylation phenotype [36, 45]. Surprisingly, *IDH2* mutations showed no statistically significant change in mtDNA abundance, suggesting that the effect is specific to the cytosolic isoform of *IDH.* Notably, mutations in *NF1,* showed a significant decrease in mtDNA abundance (BH-corrected p-value 0.03). These results echo a complementary finding by Navis and colleagues [28], who reported that a mutant *IDH1 R132H* oligodendroglioma xenograft model displayed high densities of mitochondria and increased levels of mitochondrial metabolic activity. They proposed that an increase in mitochondrial mass would increase activity of mitochondrial *IDH2* and compensate for loss of activity introduced by mutant *IDH1.*

Finally, prompted by a recent report implicating mutations in mtDNA itself with the pathology of kidney chromophobe carcinomas (KICH) [7], we investigated the connection between mtDNA copy number and mtDNA mutations in KICH. Using somatic mtDNA mutation calls provided in [7], we examined whether mtDNA-mutated samples were likely to harbor more or less mtDNA copies than unmutated samples. Surprisingly, we found that samples harboring mtDNA indels contained much higher quantities of mtDNA than unmutated samples (Mann-Whitney U-test p-value 0.002, S3). The same effect was not found when examining only single nucleotide variants. These results suggest that the presence of inactivating mtDNA mutations may induce increased mtDNA replication, perhaps as a response to inadequate mitochondrial energy production.

## 6 Discussion

The function of the human mitochondrial genome is to encode 13 proteins critical to mitochondrial respiration. In this study, we have investigated the variation of mtDNA copy number levels across many tumor types, arriving at several intriguing observations, Across more than half of the tumor types we studied, we found evidence for depletion of mtDNA, relative to normal tissues. Orthogonal measurements of transcription levels in these samples linked this variation to downregulation of mitochondrially-localized metabolic pathways, including respiration, the TCA cycle, fatty-acid oxidation, and branched-chain amino acid metabolism. Finally, we used evidence of somatic alterations in tumor samples to identify a set of mutations and copy number variants which are significantly correlated to mtDNA levels.

Our findings of gross changes in mtDNA content in tumors echo a number of prior but isolated observations, largely based on quantitative PCR measurements and with substantially smaller sample sizes, of mtDNA copy number changes in cancers (see review by [46] for a complete discussion). These include a decrease in mtDNA in breast cancer [10, 25], liver cancer [22], and clear-cell kidney cancers [27, 29]. While the majority of our observations agree with prior work (when comparing to [46]), some of our results are in contradiction to prior studies. The discordance between findings seems in part due to inadequate sample sizes, and incomplete or unavailable matched normal tissue. For example, in contrast to [25] and [43], we find no clear increase or decrease in mtDNA content in thyroid or endometrial carcinomas, respectively. However, [25] profiled 20 paired thyroid tumors, versus 116 paired thyroid tumors in this report; and [43] utilized unpaired samples of tumor and normal endometrial tissue [43], versus 41 paired samples here.

Mitochondrial DNA copy number itself reflects the cell’s capacity to conduct oxidative phosphorylation, and its depletion in many cancer types mirrors the long-standing observation that tumors reduce respiratory activity in favor of glycolytic flux *(i.e.* the Warburg effect). An interesting prospect for future study will be to identify the source of depletion of mtDNA in the cancer types like bladder and clear-cell renal cell carcinoma, where nearly all samples are depleted of mtDNA. A number of factors controlling mitochondrial biogenesis are known to be either activated *(e.g.* HIF1*α*, PGC1*α* [21]) or somatically mutated in cancers, *(e.g.* NRF2 and its regulator KEAP1 [33]) in cancers, with potential implications for mtDNA copy number.

While mtDNA depletion or accumulation may typify certain cancer types, we further identified that subsets of patients, characterized by the presence of particular somatic mutations/copy number alterations, were themselves enriched/depleted of mtDNA. The presence of activating IDH1 mutations (in low grade gliomas) or a large number of copy number alterations (in serous-like endometrial carcinomas) strongly predisposed patients to high tumor mtDNA content. If these tumors (and others with increased mtDNA content) harbor an increased dependence on mitochondrial metabolism to proliferate, this may offer a novel route for targeted therapy.

## 7 Methods

### 7.1 Data Acquisition

Whole exome sequencing (WXS) and whole genome sequencing (WGS) BAM files for 21 distinct TCGA studies were obtained from the TCGA CGHub repository (Figure 2) [44]. In total, we analyzed 13411 samples. We restricted our analyses to sequence data aligned to GRCH37 using the mitochondrial Cambridge Reference Sequence (CRS). We focused only on primary tumor, adjacent normal tissue, and normal blood samples (“01”, “11”, and “10” in the sample type field of the TCGA barcode). We further restricted our analyses to samples which were not whole-genome amplified prior to sequencing *(i.e.* we only used samples containing ‘D’ in the analyte field of the TCGA barcode), because such amplification could potentially bias the relative abundances of mitochondrial and nuclear DNA in the sample. Samtools [23] was used to extract a BAM file containing properly paired reads with minimal mapping quality aligning to the mitochondrial genome (using -F 256 -q 30 flags). The number of properly paired reads aligning to the nuclear genome was obtained by using the samtools flagstat command, and subtracting the number of properly paired reads aligning to the mitochondrial genome.

### 7.2 Purity and Ploidy Calculation and Correction

Affymetrix SNP6 arrays for tumor and normal samples were acquired for 21 cancer types from the TCGA. Arrays for each individual cancer type were processed together, quantile-normalized and median polished with Affymetrix power tools using the birdseed algorithm to obtain allele-specific intensities. PennCNV [42] was used to generate log R ratio and B-allele frequencies for each tumor. ASCAT [38] was used to generate allele-specific copy number and estimate tumor ploidy and purity using matched arrays from tumor and normal tissue.

In order to estimate mtDNA copy number, we compare the number of reads aligning to the mitochondrial genome to the number of reads aligning to a genome of known ploidy. For samples of normal tissue, we assume this known ploidy (*R*_Normal_) is equal to 2. For tumor tissue which may be infilitrated by stromal/immune cells and copy-number altered, we need to estimate the “effective ploidy” of the sample. We define this effective ploidy to be equal to

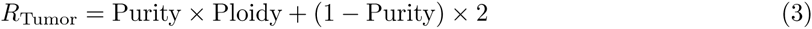

### 7.3 Survival Analysis

Survival analysis was performed with univariate Cox proportional hazards regression models where the independent and dependent variables were the log10-transformed mtDNA copy number and the overall survival respectively. The p-values for the significance of mtDNA copy number as a predictor of survival were obtained from Wald tests.

### 7.4 Gene Set Analysis

RSEM normalized RNA-Seq gene expression data was downloaded from the Broad Firehose, using the most recent data as of November 4, 2014. Data were filtered to remove genes with average read count less than 16. We calculated the nonparamemtric (Spearman) correlation (and associated p-value) between the expression of each gene and the copy number of mtDNA in the corresponding sample. To remove putatively spurious correlations, we identified all genes with a p-value less than 0.05, and set the correlation coefficient for those genes equal to zero. We then use a rank-based Mann-Whitney U-test (implemented in the geneSetTest function in the limma [20] package) to test whether particular gene sets showed an enrichment for over-or under-expressed genes, relative to the distribution of correlations across all genes. Thus, a single p-value is calculated for both alternative hypotheses 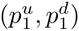. All p-values were adjusted using the Benjamini-Hochberg procedure. The method described above was applied to every Reactome pathway in the MSigDB canonical pathways gene set.

The analysis was run separately for tumor and normal tissues, and for samples processed by different sequencing center. We applied our gene set analysis pipeline to all studies for which we had at least 20 samples of RNA-Seq data (in order to retain sufficient statistical power). Analyses were run for each combination of tumor type, tissue, and sequencing center, and ensuing results were then aggregated across all studies. All results from the analyses are provided in the Supplementary Table 3.

### 7.5 Mutation and Copy Number Alteration Analysis

For each study, Gistic2 and MutSigCV results were downloaded from the Broad Firehose (most recent data as of Nov 14, 2014). From Gistic, we retained all arm-level and focal alterations with q-value less than 0.1. For mutations, we obtained the MAF file from the output of MutSig. For each gene, we calculated the number of patients in which this gene exhibited a nonsynonymous, coding mutation *(i.e.,* missense, non-sense, frameshift, in-frame insertion/deletions,and splice-site mutations), excluding those with greater than 600 non-synonnymous coding mutations). We then retained any genes which were mutated in greater than 4 percent of patients. We used only WXS data, and did not analyze tumor types which were sequenced at more than one sequencing center. Non-parametric Mann-Whitney U-test were used to evaluate whether tumors bearing a particular somatic alteration contained significantly higher/lower amounts of mtDNA in tumor samples. After testing all associations, p-values obtained from the U-tests were corrected using the Benjamini-Hochberg procedure.

## 8 Supplementary Tables

**Table 1.** Correlation of mtDNA estimates between WXS and WGS. N is the number of matched samples. Correlation corresponds to the Spearman correlation coefficient, and P-Value corresponds to the p-value of this correlation coefficient. Center: BI, Broad Insitute; BCM, Baylor College of Medicine; WUGSC, Washington University; HMS: Harvard Medical School.

| Center | Correlation | N | P-Value | Study |
| --- | --- | --- | --- | --- |
| WXS:BI::WGS:HMS-RK | 0.49 | 171 | 1.7e-11 | BLCA |
| WXS:WUGSC::WGS:WUGSC | 0.55 | 73 | 5.3e-07 | BRCA |
| WXS:BCM::WGS:BCM | 0.87 | 21 | 3.5e-07 | KIRC |
| WXS:BI::WGS:BCM | 0.48 | 25 | 0.015 | KIRC |
| WXS:BI::WGS:BI | 1 | 4 | 0 | KIRC |
| WXS:BI::WGS:BI | 0.71 | 59 | 4.5e-10 | GBM |
| WXS:BI::WGS:BCM | 0.42 | 43 | 0.0049 | HNSC |
| WXS:BI::WGS:BI | 0.3 | 34 | 0.08 | HNSC |
| WXS:BI::WGS:HMS-RK | 0.55 | 182 | 9.3e-16 | HNSC |
| WXS:BI::WGS:BI | 0.38 | 32 | 0.033 | LUAD |
| WXS:BI::WGS:HMS-RK | 0.56 | 162 | 1.2e-14 | LUAD |
| WXS:BI::WGS:BI | 0.89 | 7 | 0.0068 | OV |
| WXS:BI::WGS:BI | 0.51 | 69 | 9.2e-06 | THCA |
| WXS:BI::WGS:HMS-RK | 0.64 | 151 | 1.3e-18 | THCA |
| WXS:WUGSC::WGS:HMS-RK | 0.81 | 104 | 1e-25 | UCEC |
| WXS:WUGSC::WGS:WUGSC | 0.64 | 37 | 2.4e-05 | UCEC |

**Figure S1.**
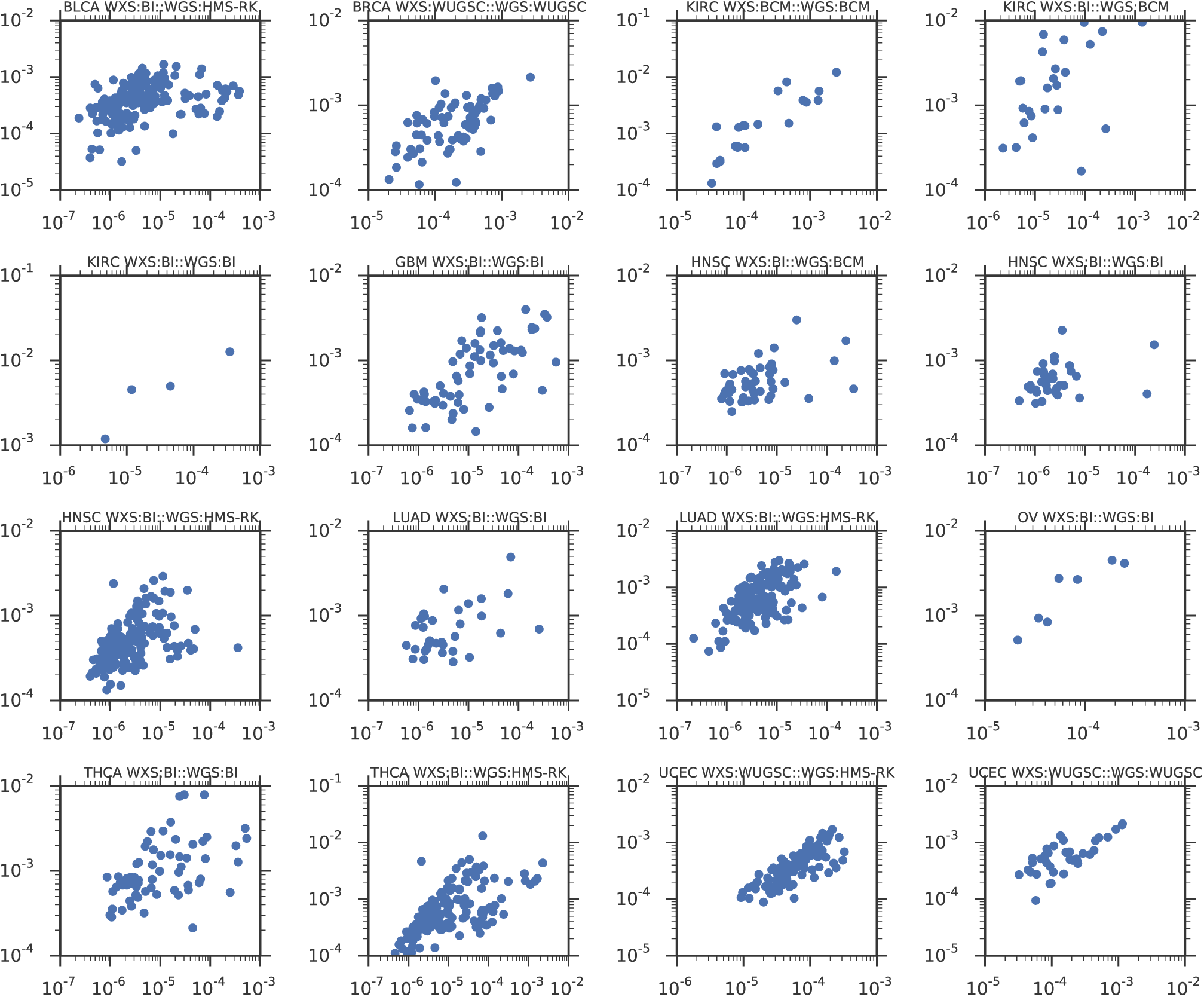
Comparison of mtDNA copy number estimates of samples profiled by both whole genome (WGS) and whole exome (WXS) sequencing. Data are separated by sequencing center, indicated in the title of each subplot.Center: BI, Broad Insitute; BCM, Baylor College of Medicine; WUGSC, Washington University; HMS: Harvard Medical School.

**Figure S2.**
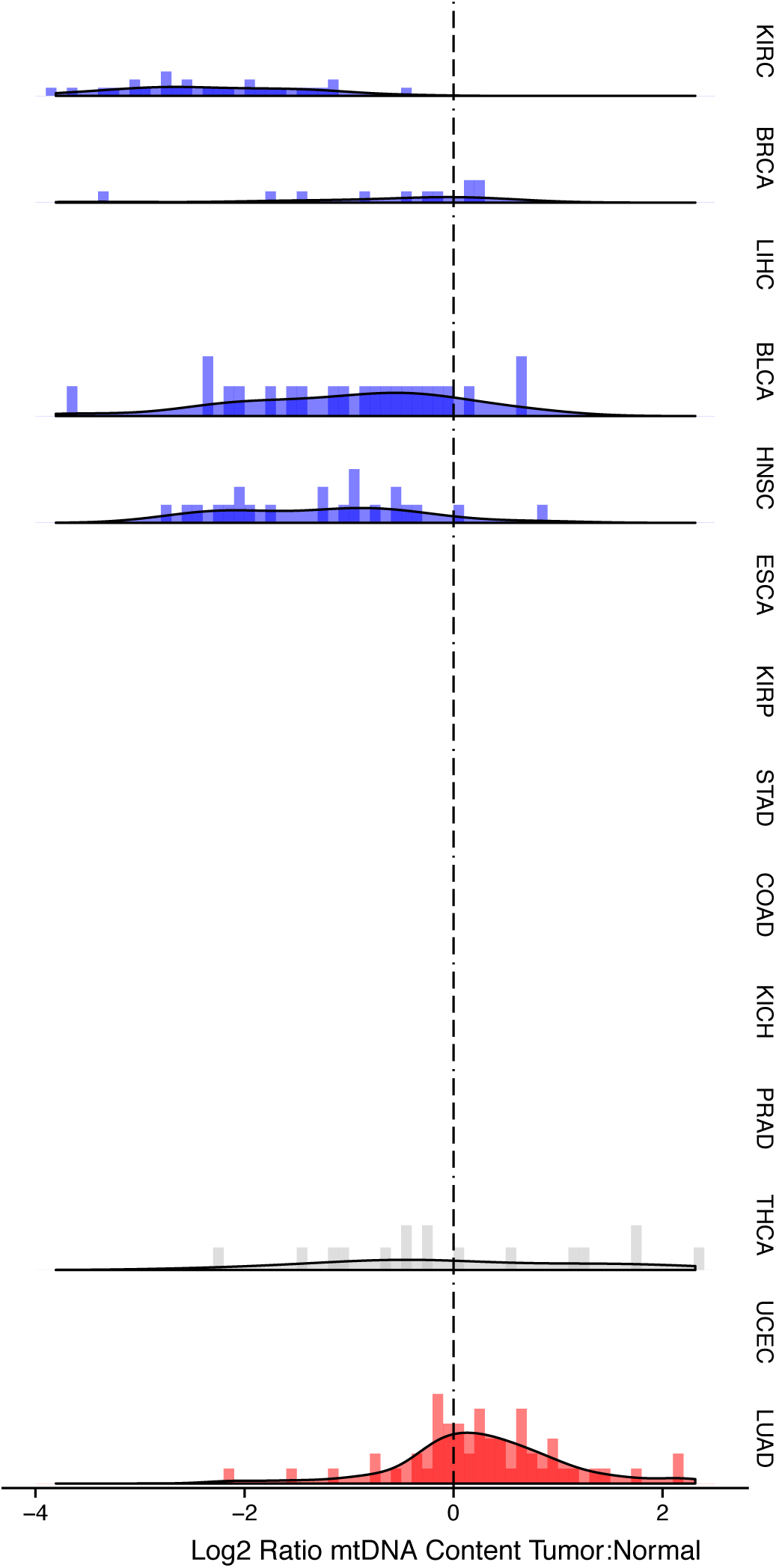
mtDNA tumor:normal copy number ratio using whole genome sequencing (WGS) data. Trends in WGS are in agreement with those in Figure 2 in whole exome sequencing (WXS).

**Figure S3.**
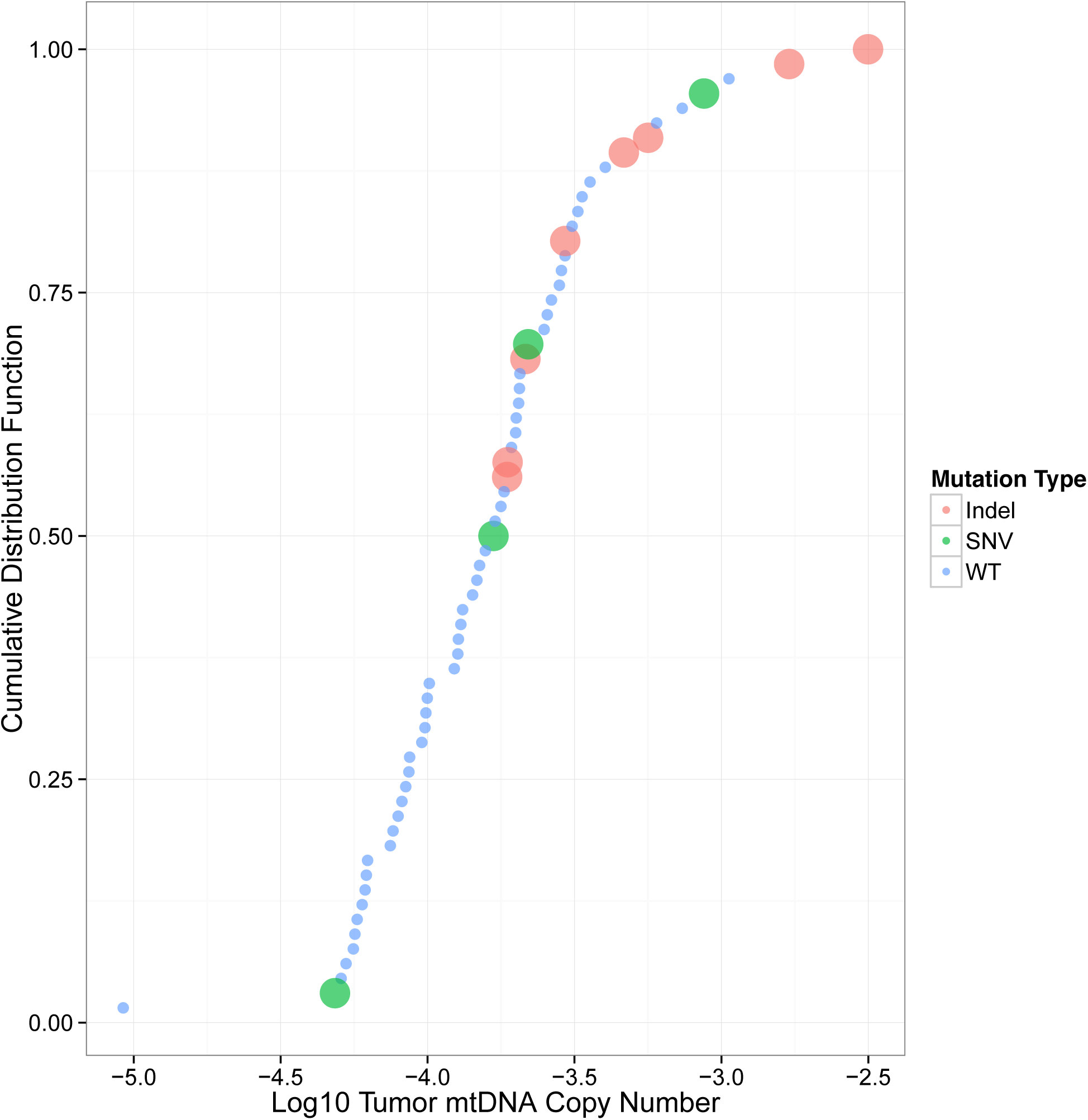
mtDNA copy number levels in kidney chromophobe carcinoma are elevated in samples with truncating mtDNA mutations. Samples are sorted by increasing mtDNA copy number on the X-axis, with percentile indicated on the Y-axis.

